# Characterization of sinking and suspended microeukaryotic communities in spring Oyashio waters

**DOI:** 10.1101/2023.09.13.557659

**Authors:** Qingwei Yang, Yanhui Yang, Jun Xia, Hideki Fukuda, Yusuke Okazaki, Toshi Nagata, Hiroyuki Ogata, Hisashi Endo

## Abstract

Microeukaryotes are important components of sinking particles contributing to carbon export from the surface to deep oceans. The knowledge of the sinking microeukaryotic communities and their dynamics is limited. We applied 18S rDNA metabarcoding method to investigate the microeukaryotic communities in sinking and suspended particles distinguished by marine snow catchers (MSC) during spring in the Oyashio region. Sinking particles displayed distinct communities and lower diversity than did suspended particles. The community compositions of the sinking particles varied with depth, suggesting that microeukaryotes were selectively removed through disaggregation or decomposition during settling. Prymnesiophyceae and diatoms were effectively removed, as indicated by their decreased abundance in the sinking particles at increasing depths. Conversely, phototrophic dinoflagellates maintained a higher abundance in the sinking particles across depths, indicating resistance to disaggregation and decomposition. Heterotrophic dinoflagellates and Spirotrichea were enriched in sinking particles and marine stramenopiles (MAST) groups were enriched in suspended particles. Sinking heterotrophic protist communities in the deep layers were similar to those in the surface layers, whereas they differed from the suspended ones in the same layer. Therefore, heterotrophic protists in surface layers were transported to deeper layers. Overall, our results demonstrate the functional differences among microeukaryotes in the biological carbon pump.

## Introduction

The sinking of particulate organic carbon (POC) is a major driver of the oceanic biological carbon pump (BCP), a process by which biologically fixed carbon is transferred from the euphotic layer to the deep ocean (Boyd *et al*., 2019). Global estimates of the sinking POC flux out of the surface mixed layer range from 5 to 20 Gt C yr^−1^ (Zhang *et al*., 2018); this influences the air-sea exchange of carbon dioxide and the Earth’s climate (Henson *et al*., 2022). Among the various types of particulate constituents of sinking POC, microeukaryotes, including photosynthetic and heterotrophic protists, play an important role in determining the structure, sinking velocity, and chemical composition of sinking POC, thereby influencing the magnitude and efficiency of the BCP (Guidi et al., 2009; Trudnowska et al., 2021). Therefore, examining the taxonomic composition and variability of the microeukaryotic assemblages in sinking POC is important.

Conventionally, microeukaryotes in sinking particles were collected using sediment traps and examined using light microscopy and biomarker pigment analysis (Scharek *et al*., 1999; Agusti *et al*., 2015). Silicified or calcified phytoplankton, such as diatoms and coccolithophorids, were often abundant in sinking particles, suggesting that their mineral cell walls (frustules or plates) function as ballasts to enhance the settlement of aggregates containing these cells or their remnants (Bauerfeind *et al*., 2009). However, these techniques have limited taxonomic resolutions and difficulty in identifying protists lacking mineral cell walls, especially when small and non-pigmented (heterotrophic). Recent advances in environmental DNA-based approaches, such as 18S rDNA metabarcoding method (Duret *et al*., 2020; Durkin *et al*., 2022), have provided comprehensive information on community composition, improving our understanding of the role of microeukaryotes in the export of carbon to the deep ocean. Small and non-mineralizing species, such as non-calcifying haptophytes and dinoflagellates, could have been underestimated in previous studies (Richardson, 2019; Lin *et al*., 2022). Time-series sediment trap studies at abyssal depths have documented the occurrence of various eukaryotic lineages of protists originating from surface layers (Boeuf et al., 2019; Preston et al., 2020).

Despite the growing knowledge of microeukaryotic assemblages associated with sinking particles, some important questions remain unresolved. One such question is whether certain microeukaryotes are preferentially associated with sinking rather than suspended particles and are more efficiently transferred from the sunlit layer to the abyssal depths. The sinking POC flux is strongly attenuated through the mesopelagic layer (200−1000 m), leaving less than 10% of the exported POC available for consumption in the bathypelagic layer and abyssal sediments (Zhang *et al*., 2018). Several mechanisms have been proposed to explain this depth-dependent POC flux attenuation, including microbial degradation and solubilization (Anderson and Tang, 2010), zooplankton consumption, and fragmentation due to the turbulence created by zooplankton swimming and feeding activities (Turner, 2015). The relative importance of the different mechanisms remains unclear; however, microeukaryotic cells that are more resistant to POC flux attenuation processes (disaggregation or decomposition) could contribute more strongly to the transfer of carbon to the deeper depths, thereby enhancing the efficiency of BCP. Conversely, easily disaggregated, or degraded microeukaryotes could contribute to carbon remineralization in the mesopelagic layer. Recently, large sedimentation chambers [marine snow catcher (MSC), (Lampitt *et al*., 1993)] were used to differentiate microeukaryotic assemblages in sinking and suspended particles. The composition of microeukaryotic assemblages differed between sinking and suspended particles in the Scotia Sea (Duret et al. 2020). In the upper mesopelagic layer, a chain-forming diatoms dominated the sinking particle assemblages, whereas prymnesiophytes were abundant in suspended particles, suggesting that diatom-enriched particles are more efficient in carbon transfer to the upper mesopelagic layer than are those enriched in prymnesiophytes. This study provided valuable insights into the functional differences among microeukaryotic taxa in the regulation of BCP; the data were limited to the upper mesopelagic layer. This hampers the assessment of microeukaryotic dynamics throughout the mesopelagic water column.

In this study, we used the MSC to investigate the characteristics of the microeukaryotic community associated with sinking and suspended particles and their changes during the sedimentation through the mesopelagic water column. Samples were collected from three depths: the subsurface chlorophyll maximum (SCM, depth 11–30 m), 10 m below the pycnocline (PC, depth 65–250 m), and the bottom boundary layer (BBL, depth 289–1489 m). This study was conducted during spring blooms in the Oyashio waters off Hokkaido in the western North Pacific Ocean (Kuroda *et al*., 2019). This oceanic region is known for its strong POC flux (Takahashi et al., 2002) and high efficiency of BCP (Kawakami and Honda, 2007) during blooms. Earlier sediment trap observations suggest that diatoms in this area play a key role in transporting POC to deep oceans owing to their large size and high sedimentation rates (Kawakami and Honda, 2007; Kawakami *et al*., 2015). However, the detailed structures of sinking particles and their decomposition processes during sedimentation from the surface to deep oceans remain unexplored.

## Results

### Sequencing statistics and environmental conditions

A total of 3,913,663 18S rDNA (V4 region) paired-end reads were obtained from 32 suspended and sinking particle fractions. After quality trimming, merging, and chimera and singleton removal, 2,416 amplicon sequence variants (ASVs) were generated. Among these, 242 ASVs taxonomically assigned to multicellular organisms were excluded; in addition, 290 ASVs that were unique to BBL samples (0.3–2% per BBL sample), potentially indigenous benthic organism, were removed (Fig. S1). The remaining 1,875 protist ASVs were subjected to downstream analyses, and their relative frequencies were normalized by rarefying the smallest number of sequences per sample (33,794 paired-end reads, Fig. S2). The protist ASVs were classified into either phototrophs (686 ASVs, including mixotrophs) or heterotrophs (1,189 ASVs) based on the taxonomic categories from PR2 (Tables S1, S2, and S3). High chlorophyll *a* (Chl *a*) concentrations were detected at St3 (5.4 µg/L) and St4 (11.9 µg/L) in March, and at St4 (7.9 µg/L) in May. Relatively higher POC fluxes (153.4–755.7 mg C m^−2^ d^−1^) and PON fluxes (30.9–129.7 mg N m^−2^ d^−1^) were observed in these sites than in the other sites (89.7–385.3 mg C m^−2^ d^−1^ and 18.4–64.8 mg N m^−2^ d^−1^) (Table S4).

### Microeukaryote community and its relationship with environmental variables

Diatoms (Bacillariophyta) were the most abundant group at St3 and St4 in March (50.5% ± 15.0% of the protists 18S reads in suspended particles at SCM; Figs. 1-2). Considering the high Chl *a* concentrations (> 5 µg/L) and high relative proportions of diatoms, we defined the St3 and St4 samples in March as “diatom-dominant” samples (Kuroda *et al*., 2019). At the genus level (level 7 category of the PR2 database), two chain-forming diatoms, *Porosira* and *Odontella*, were the most abundant at SCM (26.0% ± 8.6% and 13.0% ± 4.5% in suspended particles, 32% ± 7.5% and 10.3% ± 3.4% in sinking particles, respectively). Both genera have been previously classified as the spring-type diatoms in the Oyashio region (Chiba et al., 2004). *Porosira* and *Odontella* consistently exhibited the higher abundance at BBL in both suspended (24.2% ± 0% and 18.0% ± 1.3%, respectively) and sinking particles (16.8% ± 10.5% and 4.8% ± 1.3% respectively), even at >1000 m depth (Fig. 2). In other samples (St2 in March, and St1, 2, and 4 in May), the relative abundance of dinoflagellates was relatively higher (24.6% ± 9.2%) than that in the diatom-dominant samples (10.6% ± 4.7%); these samples are hereafter referred to as “dinoflagellate-abundant” samples. At St4, a substantial increase in the relative abundance of dinoflagellates was observed from March (6.2% of suspended particles in the SCM) to May (32.1%). Dinoflagellates constituted the group of protists with the highest relative abundance in sinking microeukaryotic communities throughout the water column in May, which showed a peak in the POC/PON fluxes (Figs. 2 and 3a; Table S1). In other stations, prymnesiophytes such as *Phaeocystis* were the abundant phytoplankton in the SCM layer (22.4% ± 4.0% in suspended particles), and they accounted for some proportions in the sinking particles at depth (2.8% ± 1.7%). However, this was significantly lower (Wilcoxon test, *p* < 0.01) than that in the suspended particles at the same depth (10.3% ± 6.2%) (Fig. 2). The relative abundance of sinking prymnesiophytes at SCM (3.8% ± 1.7%) was higher than that at PC (3.2% ± 2.0%) and at BBL (2.5% ± 1.7%). Vector analysis using vegan’s “envfit” function showed that the surface Chl *a*, temperature, and PON flux were significantly correlated with the variation in the eukaryotic community in the suspended particles (*p* < 0.05) (Fig. 3a).

**Fig. 1.**
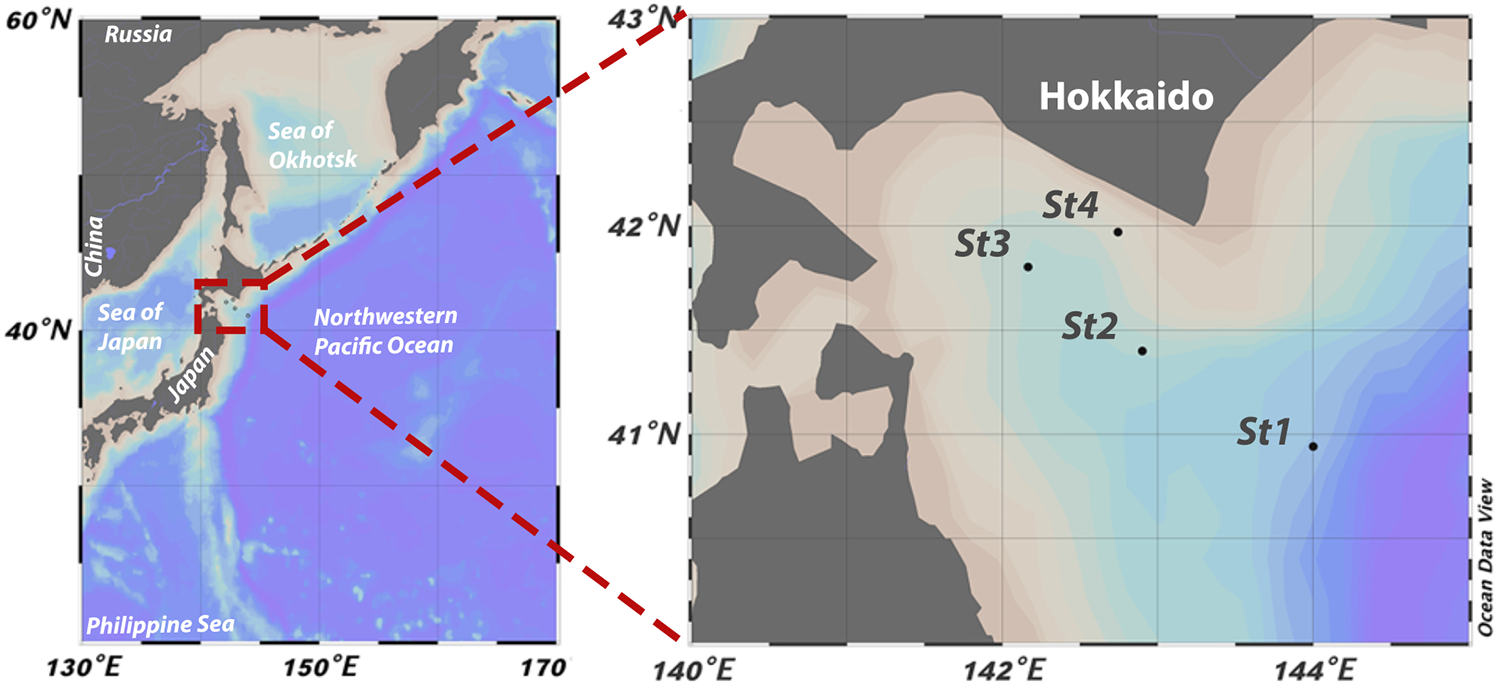
Location of sampling sites drawing by Ocean Data View.

**Fig. 2.**
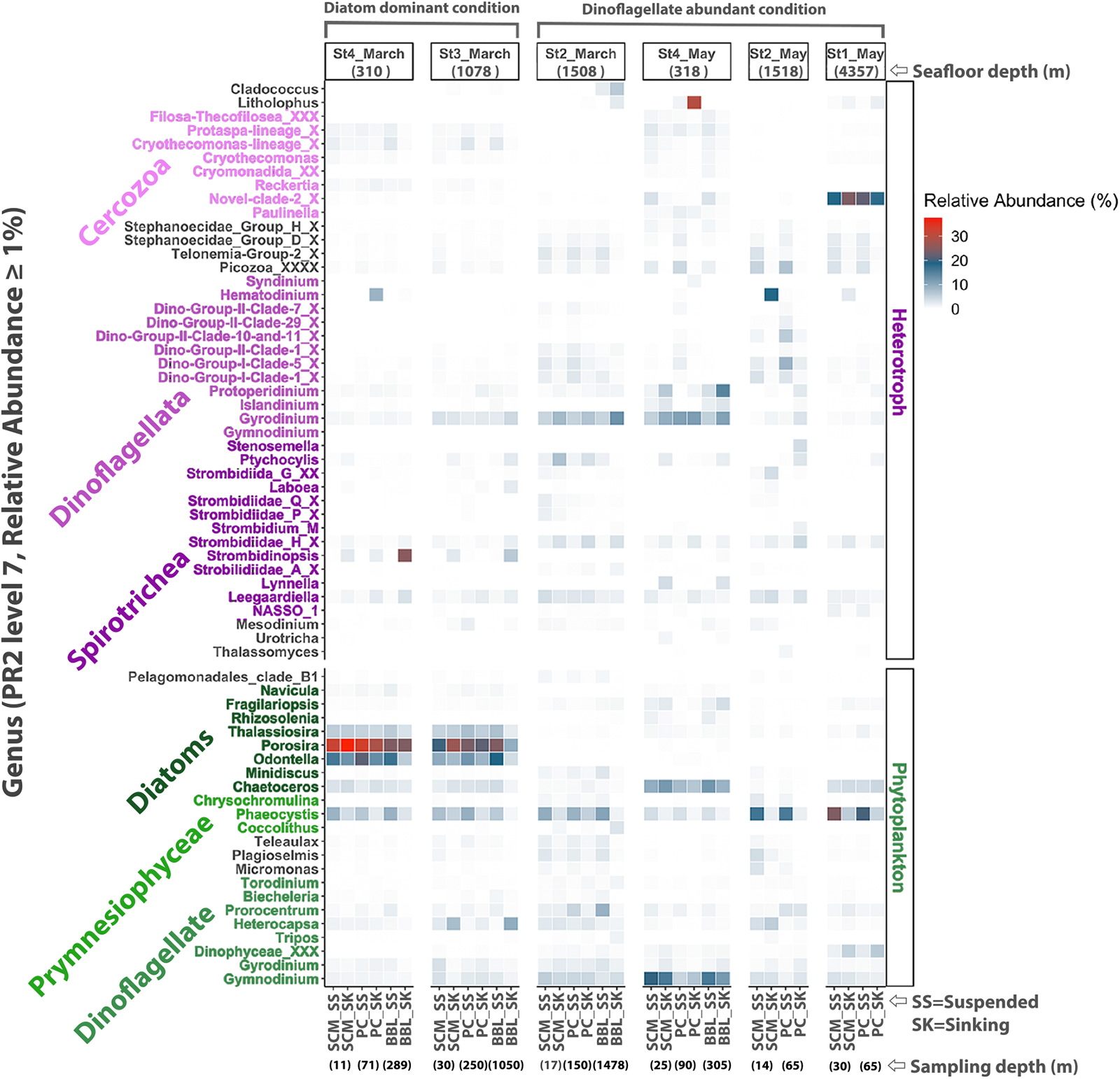
Overview of microeukaryote composition in Oyashio waters. The relative abundance of the protists genus (level 7 category of the PR2 database, relative abundance ≥ 1%), colored based on major divisions or class information at suspended and sinking particle samples across stations and depths; X-axis indicates particle fractions of each depth: SCM_SS=Suspended particle at SCM, PC_SK=Sinking Particle at PC.

**Fig. 3.**
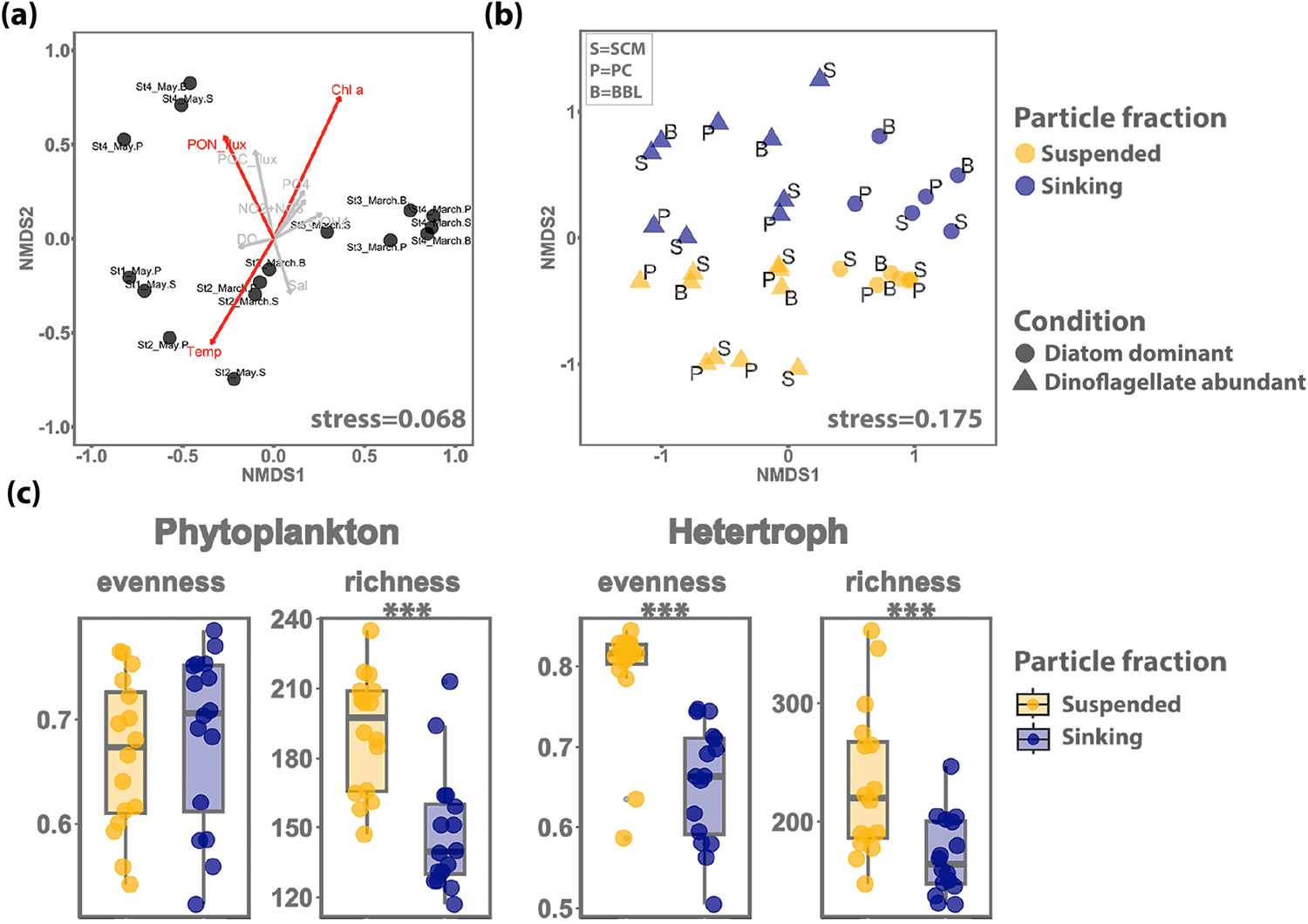
Comparison of microeukaryotic communities detected in different particle fractions. (a) NMDS plot of microeukaryote protists in suspended particles, including 10 fitted environmental variables: total nitrogen (NO2+NO3), phosphate (PO4), silicate (Si(OH)4), temperature (Temp), dissolved oxygen (DO), salinity (Sal), chlorophyll concentration (Chl a), and PON and POC fluxes.The variables with p < 0.05 were marked with red; (b) MNDS of Bray-Curtis dissimilarities of rarified read counts. Particle fractions were sinking particles (midnight blue) and suspended particles (golden yellow). Circle points represent the diatom-dominant condition, and triangle points represent the dinoflagellate-abundant condition. Letters indicate depth: S=SCM, P=PC, B=BBL; (c) Evenness and richness indices of different functional groups pooled by particle fractions. Stars indicate significant differences in indices between these two particle fractions, with * *p* < 0.05, ** *p* < 0.01, *** *p* < 0.001 (Wilcoxon test).

### Microeukaryotic communities differ between suspended and sinking fractions

Non-metric multidimensional scaling (NMDS) showed a clear and significant separation in the ASV composition of the microeukaryotic community between suspended and sinking particles (PERMANOVA, *p* < 0.05) (Fig. 3b, Table S5). Phytoplankton richness was lower in the sinking particles than in the suspended particles (Wilcoxon test, *p* < 0.05; Figs. 3c and S3). Heterotrophic protists exhibited significantly lower richness (Wilcoxon test, *p* < 0.05) and *Pielou*’s evenness (Wilcoxon test, *p* < 0.05) in the sinking particles than in the suspended particles (Fig. 3c).

Phytoplankton communities exhibited consistently high similarity index between sinking and suspended particle types with values between 0.54 and 0.63 at each depth, and between different depths (the Bray-Curtis similarity: 0.58–0.60 for sinking particles and 0.56–0.59 for suspended particles) (Figs. 4 and S4). Communities of heterotrophic protists also showed higher similarities between sinking and suspended particles at SCM (0.60 ± 0.12), but the values were lower at PC and BLL (0.40 ± 0.24 and 0.43 ± 0.21, respectively). Furthermore, the communities of sinking heterotrophic protists at depth showed higher similarity to those in the sinking particles from the upper layer (0.55–0.66) than to those in the suspended particles from the same layer (0.43–0.48) (Figs. 4b-c). Conversely, suspended heterotrophic communities at depth showed lower similarity to their upper-layer counterparts (0.39– 0.51) (Figs. 5S and 4c).

**Fig. 4.**
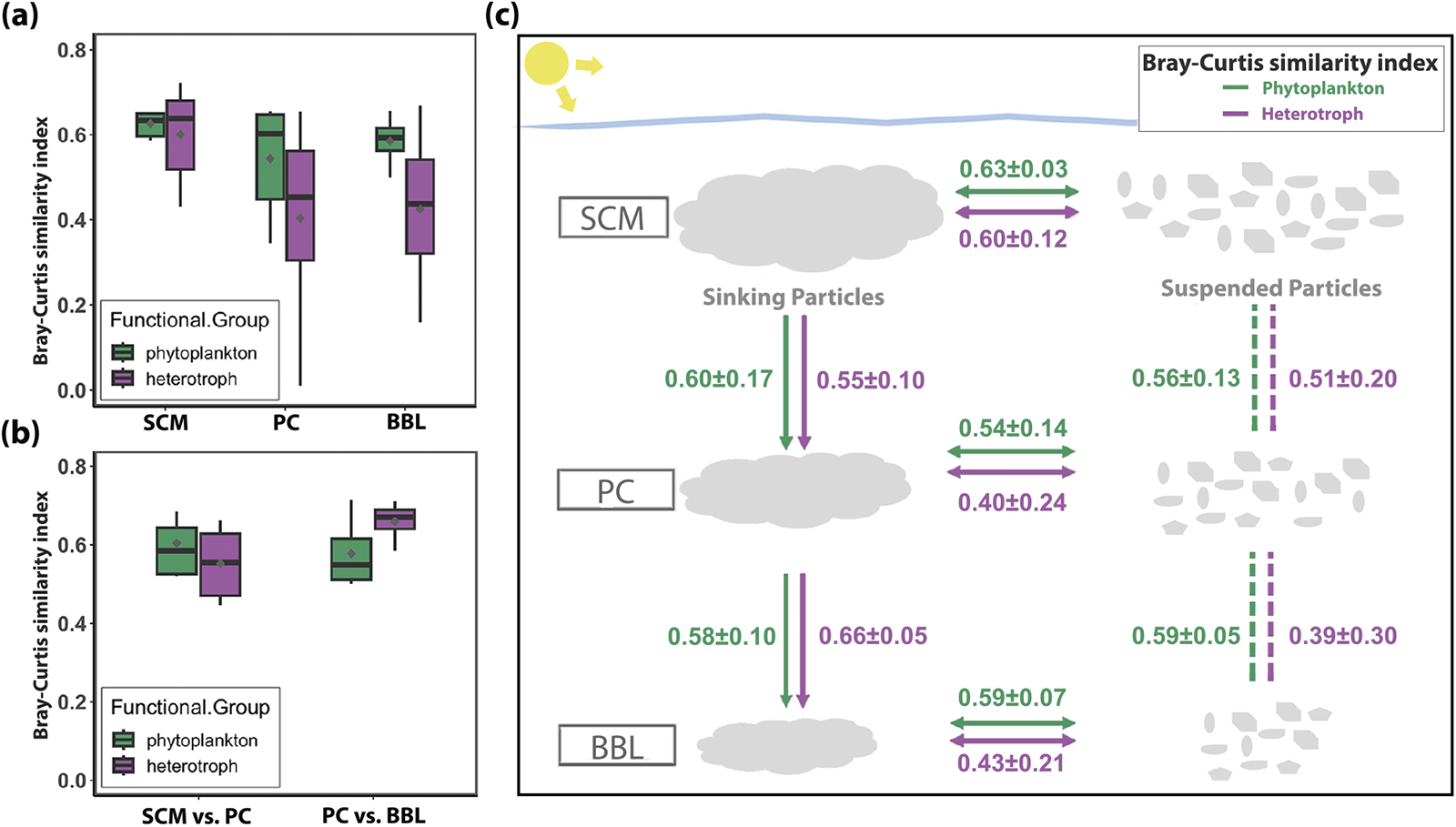
Comparison of phytoplankton and heterotrophic protists communities. (a) Boxplot showing the Bray-Curtis similarity between suspended and sinking phytoplankton and heterotrophs across depth. (b) Boxplot showing the Bray-Curtis similarity of sinking phytoplankton and heterotrophic communities between different depths. Grey points in the plots represent the mean value of the observations. Outliers were removed from the boxplots. (c) Comparative schematic diagram of similarity of phytoplankton (green) and heterotrophic protists (purple) communities. Values indicate the Bray-Curtis similarity index.

**Fig. 5.**
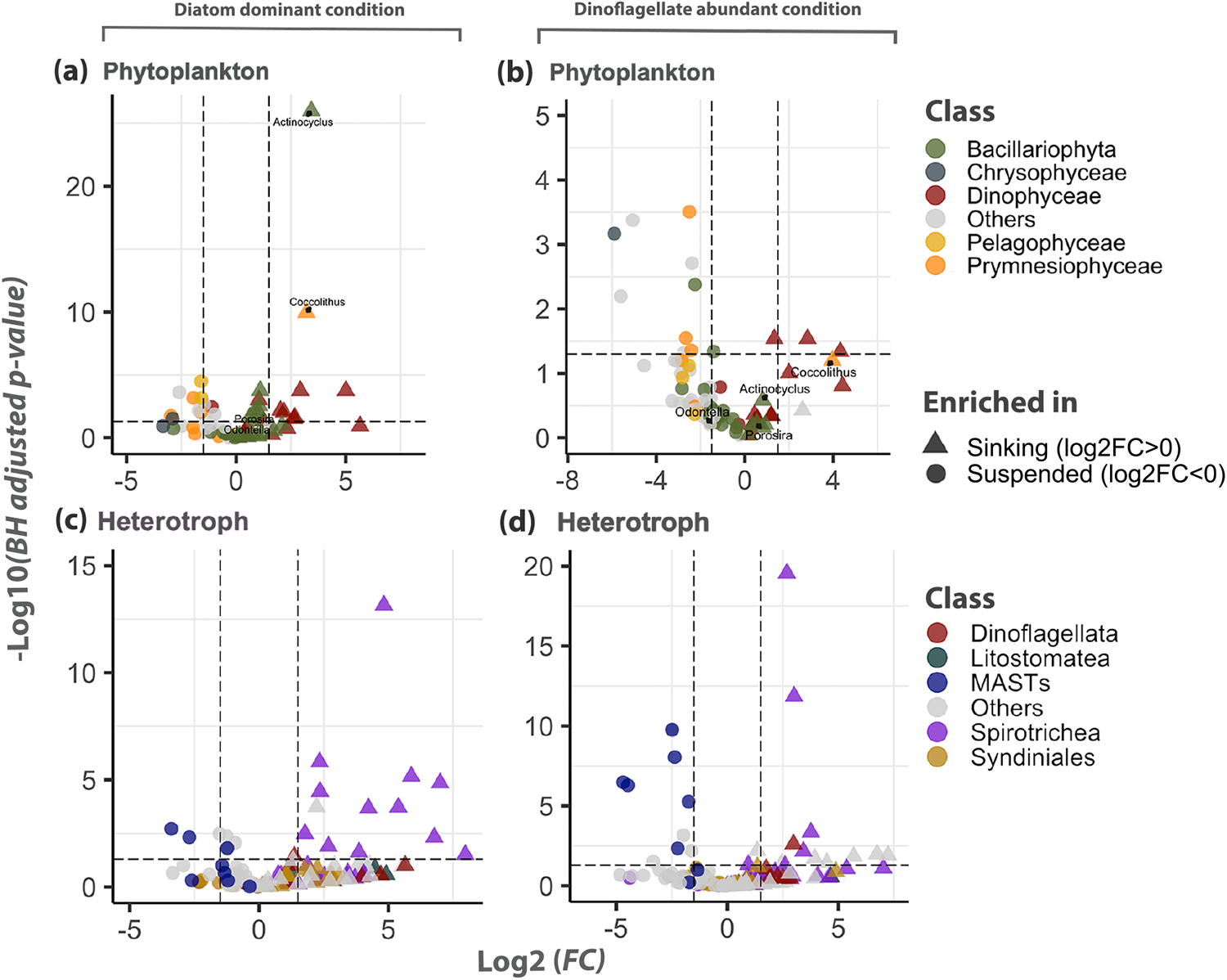
Differential abundance of functional groups at different conditions. The volcano plot depicts the differential abundance of phytoplankton and heterotrophs at diatom dominant condition (a&c) and dinoflagellate abundant condition (b&d). Log2 (fold change) (FC) is plotted against -log10 (BH adjusted p-value) mostly at the genus level. Each dot represents a genus, color-coded based on their division or class information. For those unassigned at the genus level, higher taxonomic information was retained up to the family level. Taxa unassigned at the higher than family level was removed. Significantly distinct genera between suspended and sinking particles were defined based on the absolute of log2 (FC) > 1.5 and BH adjusted *p*-value < 0.05. Log2 (FC) > 0 represents taxa enriched in sinking particles (triangle), while log2 (FC) < 0 indicates taxa enriched in suspended particles (circle).

The NMDS results also showed that the diatom-dominant samples were significantly different from the dinoflagellate-abundant samples (PERMANOVA, *p* < 0.001; Fig. 3b, Table S5). Differential abundance analyses revealed that the protist genera showed varying patterns in relative abundance between the suspended and sinking particles in the samples under both conditions (Fig. 5). Under diatom-dominant conditions, most of the diatom genera (except *Actinocyclus*), including the dominant *Porosira* and *Odontella*, displayed no significant difference between the suspended and sinking particle fractions (Fig. 5a). Most phototrophic dinoflagellates (16 out of 18 genera) were more abundant in the sinking particles than in suspended particles, with five genera showing significant differential abundances (Benjamini– Hochberg (BH) adjusted *p*-value <0.05). Most genera belonging to Prymnesiophyceae (except *Coccolithus*) (7 out of 9 genera), Chrysophyceae (2 out of 2 genera) and Pelagophyceae (4 out of 4 genera) were more enriched in the suspended particles than in sinking particles (Fig. 5a), with five genera of which showed statistically significant differences (BH adjusted *p*-value < 0.05). These trends in phytoplankton distribution across fractions were also evident under dinoflagellate-abundant conditions (Fig. 5b). Regarding the heterotrophic protists in the diatom-dominant condition, several Spirotrichea (a class of ciliates) genera and heterotrophic dinoflagellate genera were significantly (BH adjusted *p*-value < 0.05) enriched in the sinking particles (Fig. 5c). In contrast, the MAST lineages (MAST-1A, MAST-2, and MAST-7), which contain heterotrophic nanoflagellates, were primarily found in the suspended particles (BH adjusted *p*-value < 0.05). Similar to phytoplankton communities, heterotrophic protists showed divergent distribution patterns between the suspended and sinking fractions in the dinoflagellate-abundant samples (Fig. 5d).

### Depth-dependent attenuation of sinking particle-associated microeukaryotes

The composition of key protist classes (level 4 category of the PR2 database) changed with depth in the sinking particles. The relative abundance of phototrophs decreased (from 64.8% ± 14.0% at SCM to 48.6% ± 2.2% at BBL), while that of heterotrophs increased with depth (Fig. 6a). In particular, the dominant *Porosira* and *Odontella* genera displayed remarkable decreases in relative abundance with depth, at St4 and St3 in March (Fig. 6b). Conversely, the relative abundances of Spirotrichea and heterotrophic dinoflagellate increased with depth (Fig. 6a). This depth-dependent trend was more pronounced for the relative abundances at the family level (level 5 category of the PR2 database) (Fig. 6c).

**Fig. 6.**
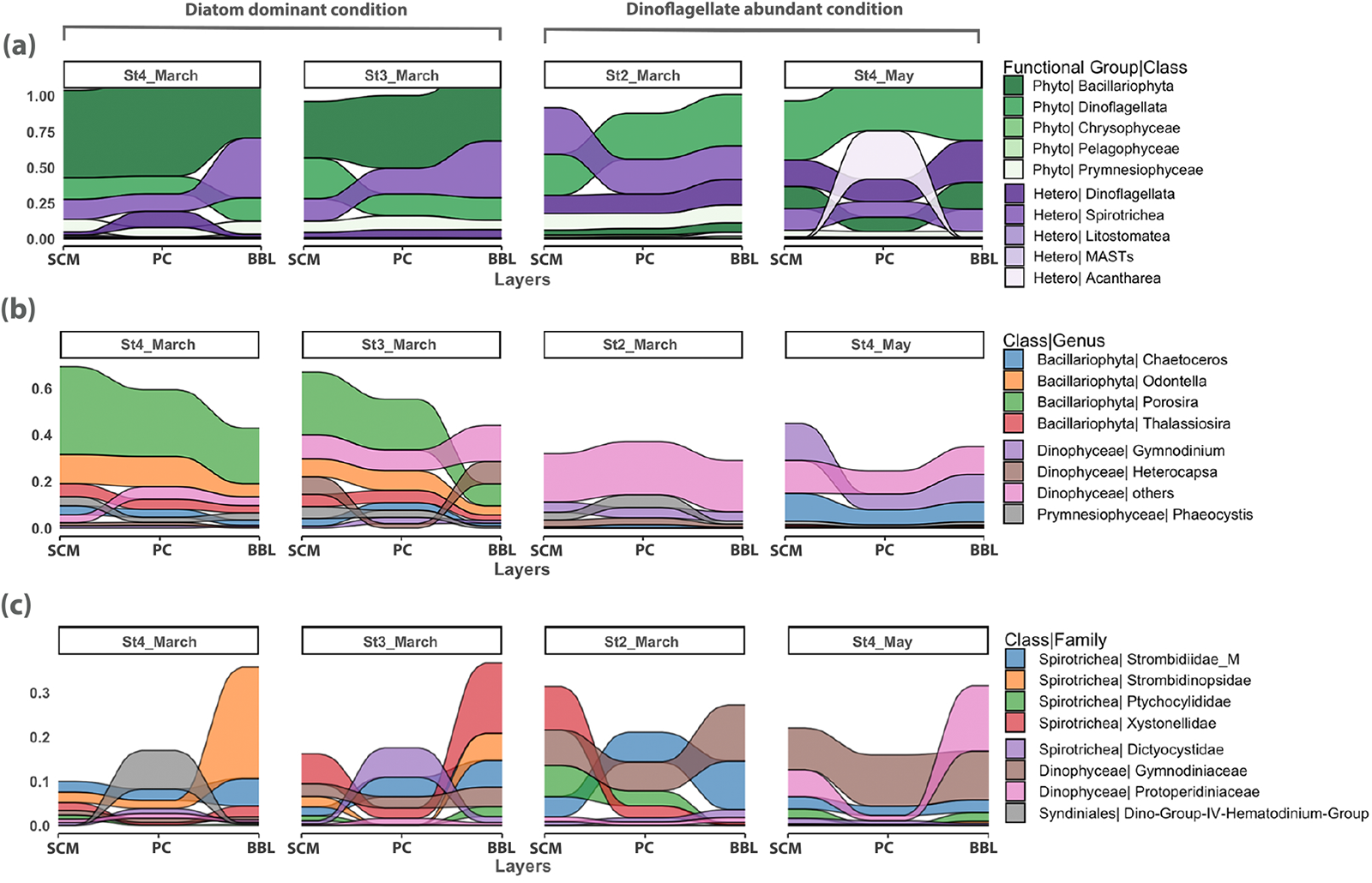
The vertical succession of sinking particle-associated (a) major microeukaryotes composition at the class level, then further separated by phytoplankton at the genus level (relative abundance ≥ 5%) (b) and heterotrophic protists at the family level (relative abundance ≥ 5%) (c) Stations without bottom layers (St1 and St2 in May) were removed from the analysis.

## Discussion

We observed a clear separation of phytoplankton composition between the suspended and sinking particles (Figs. 3b–c and 5a–b). The lower phytoplankton richness in the sinking particles, particularly in the SCM (Fig. S3), could be attributed to selective aggregation and sinking processes. Namely, a subset of the microeukaryotic taxa in the suspended particles is incorporated into the aggregates; this contributes to the sinking processes. The selectivity for aggregation and sinking could be influenced by the physical and physiological properties of the phytoplankton (size, shape, stickiness, and secretion of exopolymers), which can affect the aggregate formation and sedimentation (Jackson, 1990).

Prymnesiophytes and pelagophytes, belonging to pico- and nanoplankton (0.2–20 µm), were enriched more in the suspended particles (Figs. 2 and 5a-b), possibly due to their small cell size (Liu et al., 2009; Wetherbee et al., 2015). These picophytoplankton groups were involved in sinking particles in the surface layer; however, they were rarely detected in the sinking particles in the deep layer, indicating that they were more prone to disaggregation or decomposition during the sinking process. This observation is consistent with that of a previous study showing that prymnesiophyte-enriched particles exported from the euphotic zone are likely to disintegrate into suspended particles in the upper mesopelagic layer (Duret et al., 2020).

Phototrophic dinoflagellates, which were more abundant in sinking particles and showed stable relative abundance with depth (Figs. 5a-b and 6a-b), play a significant role in the BCP across various ecosystems (Guidi *et al*., 2016; Durkin *et al*., 2022; Lin *et al*., 2022); Dinoflagellates are more carbon-dense than diatoms, as evident from their carbon-to-cell volume ratio (Menden-Deuer and Lessard, 2000); their sinking can sequester more carbon per volume of particles than that of diatoms. The formation of massive carbon-rich and thick-walled resting cysts during dinoflagellate blooms makes them more resistant to degradation, contributing to the intense sedimentation of organic matter (Heiskanen, 1993; Cohen *et al*., 2021). Thus, phototrophic dinoflagellates can act as important contributors to BCP in Oyashio waters. This is supported by our results, which show a positive correlation between dinoflagellate abundance and POC/PON fluxes (Fig. 3a).

Diatoms play a key role in POC sedimentation in the Oyashio waters, as suggested by previous studies (Kawakami and Honda, 2007; Kawakami et al., 2015). In this study, centric diatom genera such as *Porosira* and *Odontella* were dominant in the diatom-blooming sites near the coastal area (Figs. 1 and 2). These genera belong to microphytoplanktons, which have relatively large cells (20–200 µm) (Olenina *et al*., 2006). Diatoms are typically considered important contributors to BCP in various ecosystems, as heavy silica shells serve as ballast for marine snow and fecal pellets (Belcher *et al*., 2016; Tilstone *et al*., 2017). Previous observations of spring Oyashio water showed that *Porosira glacialis* was the most dominant species during bloom period, contributing to 25% of the diatom carbon biomass (Suzuki *et al*., 2011). The genus *Porosira* is enriched in the sinking particles at BBL (16.8 ± 10.5%, Fig. 2), indicating that the dominant diatom species would be the major contributors to vertical carbon export. However, the relative abundance of *Porosira* sharply decreased with depth, especially from PC (21.7%) to BBL (9.4%) at St3 in March (Figs. 6a-b). This indicates that the dominant diatoms displayed high attenuation within the sinking communities during sedimentation. These findings are consistent with that from previous field studies, which reported that the export efficiencies are lower in the diatom-blooming sites than in the non-blooming sites (Francois *et al*., 2002; Fischer and Karakaş, 2009; Henson *et al*., 2012). Sinking aggregates composed of diatoms exhibit higher porosity, which reduces particle density and sinking velocity despite high ballast availability, thus increasing their susceptibility to remineralization (Francois *et al*., 2002; Bach *et al*., 2019). Furthermore, the transparent exopolymer particles (TEPs) produced by diatoms (Engel, 2000), which facilitate aggregation, can reduce the sinking velocities of aggregates owing to their low density (Mari et al., 2017). Indeed, the contribution to vertical carbon export varies among diatom genera or species owing to differences in cell size, shape, degree of silicification, and production of TEPs (Tréguer *et al*., 2018).

The heterotrophic protist community structure also differed significantly between the sinking and suspended particles (Figs. 3b-c and 5c-d). Lower richness and evenness of the heterotrophic protists in the sinking particles than those in the suspended particles indicate that specific protist groups were enriched in the sinking particles. The prominent heterotrophic protist groups enriched in sinking particles were Spirotrichea and heterotrophic dinoflagellates (Figs. 5c-d). These protists are active grazers of phytoplankton, including chain-forming diatoms (Montagnes and Lessard, 1999; Sherr et al., 2013), suggesting that they contribute to the decomposition and transformation of diatom-enriched aggregates in Oyashio waters.

The MAST groups of heterotrophic protists were more abundant in the suspended than in the sinking particles. In addition, they were more abundant in the surface layers than in the deeper layers, which is consistent with previous observations (Obiol et al., 2021). The MAST groups include consumers of heterotrophic bacteria and picoeukaryotes in the water column (Lin et al., 2012; Orsi et al., 2018) and parasites of diatoms in sediments (e.g., MAST-6, (Rodríguez-Martínez *et al*., 2020)). Considering their low abundance in sinking particles, it is less likely that the MAST groups in suspended particles collected from the deeper layers of the Oyashio region were those transported from the upper layer. MAST groups prefer a free-living, suspended lifestyle and consume picoeukaryotes and bacteria. The distinct occurrences of heterotrophs in sinking and suspended particles reflects the differences in lifestyle and feeding habits (Worden *et al*., 2015).

High similarities in the sinking microeukaryote communities between the surface and deep layers were recorded; this included both phytoplankton and heterotrophic protists (Fig. 4c). Heterotrophic protist communities of sinking particles displayed more remarkable similarities across different depths (i.e., SCM, PC, and BBL) than did suspended communities in ambient waters. This suggests strong vertical connectivity of heterotrophic protist communities with sinking particles, mediated by the association of specific groups of heterotrophic protists (Spirotrichea and heterotrophic dinoflagellates).

In BBL samples collected approximately 10 m above the seafloor, the resuspension of bottom sediments, indicated by high turbidity, could introduce benthic communities from the sediments into the sinking and suspended particles collected with MSC. These benthic communities harbor highly diverse and abundant indigenous eukaryotes that are unique to the environment (Cordier et al., 2022). However, the BBL samples did not show an increased richness of microeukaryotes compared to that in the upper samples (Fig. S4). The proportion of ASVs unique to the BBL samples was consistently low (0.3–2%). These results suggest that resuspension of the benthic microeukaryotic community was minimal in the studied region. Alternatively, the resuspended microeukaryotic communities mostly originate from recently settled fresh particles (Almroth-Rosell et al., 2011; Rodil et al., 2020). In either case, our data suggested that heterotrophs in the sinking particles mainly originate from the surface layers along with the formation of sinking aggregates rather than through active colonization from ambient water.

Both the phototrophs and heterotrophs associated with sinking particles in the deeper layers were derived from the surface; they displayed distinct depth-dependent attenuation patterns (Fig. 6). The contribution of phototrophs decreased with depth, whereas that of the heterotrophic protists increased. The increased abundance of heterotrophic protists relative to phototrophs with increasing depth could reflect the consumption of phototrophs during the transit of sinking particles in the water column. Therefore, heterotrophic protists such as Spirotrichea and heterotrophic dinoflagellates could contribute to the decomposition of sinking particles by consuming phototrophic cells. The depth-dependent attenuation of the relative abundance of phototrophs was more prominent in diatom-dominated situations than in dinoflagellate-abundant situations, implying that diatoms are more rapidly consumed by heterotrophic protists. These results indicate that the magnitude of sinking POC flux attenuation is influenced by dominant phototrophic microeukaryotes (diatoms vs. dinoflagellates) and their consumers (Spirotrichea and heterotrophic dinoflagellates), which comprise the microbial consortia of sinking particles. Future studies are needed to elucidate the role of prey-predator interactions of microeukaryotes, which are potentially highly selective in feeding (Lima-Mendez et al., 2015), in regulating the POC flux attenuation with depth.

### Conclusion

The use of MSC combined with DNA metabarcoding to investigate spring Oyashio waters allowed us to characterize the microeukaryotic communities that contribute to the biological carbon sequestration processes. This study provides profound insights into the composition and dynamics of microeukaryotic communities within sinking and suspended particles in the Oyashio region of the western subarctic Pacific. We present the distinct structures of the microeukaryotic communities in different particle types and at varying depths. In addition, these findings highlight the differential patterns of depth-dependent attenuation among different taxonomic groups; prymnesiophytes and diatoms were more rapidly attenuated with depth than phototrophic dinoflagellates. We found strong vertical connectivity between some heterotrophic protists (Spirotrichea and heterotrophic dinoflagellates) associated with sinking particles, suggesting that they play a crucial role in the regulation of sinking POC fluxes through the consumption of major phototrophic microeukaryotes associated with sinking particles.

## Materials and methods

### Cruises and sampling strategy

Sampling was conducted at four stations (Fig.1) during the KS-21-4 (March 11–21, 2021) and KS-21-7 (May 3–11, 2021) cruises of the R/V *Shinsei-Maru* (JAMSTEC) in the Oyashio region off Hokkaido, Japan. Samples for DNA analysis were collected using MSCs at three depths: the subsurface chlorophyll maximum (SCM), 10 m below the pycnocline (PC), and 10 m above the bottom (BBL). Detailed information on the sampling depths and environmental factors is presented in Table S4.

Particles of suspended and sinking particles were sampled from seawater in the upper and base parts of MSC, respectively, then filtered using a 0.8 µm pore-size cellulose acetate membrane (47 mm diameter, Millipore) with a gentle vacuum (<0.013MPa). The filters were stored in 1.5-mL cryotubes previously filled with 600 µL of buffer RLT Plus (Qiagen), 6 µL of 2-mercaptoethanol (Sigma-Aldrich), and 0.2 g of glass beads; the samples were flash-frozen in liquid N_2_ and stored in a deep-freezer until analysis on land. The suspended fraction defined as the particles remaining in the upper part of the MSC, and the sinking fraction referred to the particles that sunk to the base plate of the MSC after being placed on the deck for 2 h. For detailed descriptions of MSC, refer to earlier reports (Duret *et al*., 2019; Baumas *et al*., 2021). The sinking fraction contained some suspended particles; therefore, after detaching the upper part of the MSC, the base of the MSC was allowed to settle for another 10 min, and approximately two-thirds of the seawater in the base plate of the MSC was siphoned from the top to remove as many suspended particles as possible.

### Measurements of environmental variables

Vertical profiles of temperature, dissolved oxygen (DO), salinity, Chl *a* fluorescence, photosynthetically available radiation (PAR), and turbidity were measured at each site using an SBE 911-plus CTD system (Seabird Electronics, Bellevue, WA, USA) (Fig. S1). Seawater samples for Chl *a* and macronutrient analysis were collected using Niskin bottles attached to a CTD-Carousel multiple-sampler system. Samples for Chl *a* concentration measurement were collected on GF/F filters and measured with a fluorometer (10-AU, Turner Designs) using the acidification method (Holm-Hansen *et al*., 1965). The concentrations of macronutrients (Si(OH)_4_, NO_2_, NO_3_, NH_4_, and PO_4_) were determined using a flow injection analyzer (AACSIII, Bran+Luebbe). The detection limits of the macronutrient measurements were defined as three times the deviation of the repeated measurements from the blank. Detailed procedures for Chl *a* and macronutrient analyses have been described by Fukuda *et al*. (2016). Samples for POC and PON concentrations were collected from the sinking fraction of MSC and analyzed using an elemental analyzer (Elemental Analyzer-IRMS; FLASH 2000, Thermo Fisher Scientific). The POC and PON sinking fluxes were calculated using the method described by Yamada et al. (submitted).

### DNA extraction and 18S rRNA gene amplification and sequencing

AllPrep DNA/RNA Mini Kits (Qiagen) were used for DNA extraction. The sample vials were agitated thrice at 2,500 rpm for 50 s using a homogenizer (µT-12, TAITEC) before proceeding with the manufacturer’s protocol (BBL samples at St1 and St2 in May failed to yield sufficient DNA for downstream analysis). For each sample, three PCR amplifications targeting the V4 region of the 18S rRNA gene were performed using the following primer pairs: the forward E572F (5’-CYGCGGTAATTCCAGCTC-3’) and the reverse E1009R (5’-AYGGTATCTRATCRTCTTYG-3’; 436 bp), as designed previously (Comeau et al., 2011). The PCR mixtures were prepared in a 25-µL reaction volume containing 1 x KAPA HiFi HotStart ReadyMix, 1 µM each of forward and reverse primers, and 2.5 ng of template DNA. The PCR consisted of a pre-denaturation step at 98°C for 30 s, followed by 30 cycles of denaturation at 98°C for 10 s, annealing at 61°C for 30 s, and extension at 72°C for 30 s, and a final extension at 72°C for 5 min. The triplicate PCR products were mixed after purification using Agencourt AMPure XP beads (Beckman Coulter). The quality and quantity of the amplicons were checked using the D1000 ScreenTape Assay kit read on a 2100 TapeStation system (Agilent Technologies) and a Qubit dsDNA BS assay kit (Invitrogen), respectively. Libraries were prepared using the Nextera XT index kit V2 and sequenced on an Illumina MiSeq platform using a 300 bp paired-end sequencing. All raw sequencing data are available at the DDBJ Sequence Read Archive under accession number of DRA015710.

### Analysis of sequencing data

Raw sequencing data were preprocessed using QIIME2 (Bolyen *et al*., 2019) (version 2021.11). Trimming of low-quality reads and primer sequences, merging of paired-end reads, dereplication, chimera removal, and ASV inference were performed using denoising algorithms in DADA2 (Callahan *et al*., 2016). A naïve Bayes classifier trained (Bokulich *et al*., 2018) on the PR2 database (Guillou *et al*., 2013) (v 4.14.1) with the E572F/E1009R primer set was used to assign taxonomy to the ASV sequences. The feature table of read counts and taxonomy assigned to ASVs was exported by eliminating singleton ASVs. The ASVs that appeared only at the BBL samples were removed to reduce the potential contamination by indigenous benthic organisms.

Potential multicellular eukaryotic ASVs, such as Embryophyceae, Fungi, Metazoa, and Rhodophyta, were excluded; the unicellular eukaryotic (protists) ASV taxa were divided into photosynthetic and heterotrophic groups (Table S3) (Simon *et al*., 2009; Durkin *et al*., 2022). We manually controlled the detailed trophic groups of dinoflagellates based on existing literature; however, some of the heterotrophic dinoflagellate sequences could have been erroneously assigned to photosynthetic groups (Table S4) or vice versa. In addition, owing to the copy number variation in the 18S rRNA genes, the relative abundance of the ribosomal marker genes does not correlate with cell abundance (Zhu et al., 2005). This potentially affects the microeukaryote composition analysis results; however, it does not affect the comparison of specific taxa between the suspended and sinking particle fractions.

### ASV-based diversity analyses

Statistical analyses were performed using R v.4.4.1 and Python v.3.9.13. Samples were rarefied to account for differences in library size, and rarefaction curves were used to investigate the degree of sample saturation by calling the ‘rrarefy’ function in the vegan v.2.6.4 package (Oksanen et al., 2022). NMDS analysis of the dissimilarity in the microeukaryotic communities between these two particle fractions, and the relationship between the suspended microeukaryote composition and environmental variables (NH_4_ was omitted because 11 out of 16 samples had concentrations below the detection limit, Table S4) were performed based on the Bray-Curtis dissimilarities with the rarefied abundance table. ANOSIM with 9999 permutations was used to statistically test the significant dissimilarity between the particle fractions (with the it‘envfit’ function in vegan). Alpha diversity indices (*Pielou’*s evenness and richness) of phytoplankton and heterotrophs in each sample were evaluated by rarefying the read numbers. The difference in alpha diversity between these two particle fractions was assessed using the Wilcoxon signed-rank exact test to determine statistical significance.

### Taxon-based differential abundance analysis

Differential analysis of protist genera between the two particle fractions was performed using DEseq2 v1.38.3 (Love et al., 2014). The samples were categorized into two conditions based on the NMDS analysis (Fig. 3b): diatom-dominant regions, including St3 and St4 in March, and dinoflagellate-abundant regions with a relatively higher abundance of dinoflagellates, including St2 in March and St1, St2, and St4 in May. Protist genera with log_2_(fold change) (FC) > 1.5 or < −1.5 and an adjusted *p*-value < 0.05 determined using the Benjamini-Hochberg method were regarded as “significant”.

## Acknowledgments

We gratefully acknowledge the captain, officers, and crew of the R/V Shinsei-Maru for their great assistance during the cruise. We are thankful to Junyi Wu and Ruixuan Zhang for support with data analysis and helpful discussions. This study was funded by JSPS KAKENHI (nos. 19H05667, 22H00384, and 22H02420). Computational work was performed at the SuperComputer System, Institute for Chemical Research, Kyoto University.

