## Supplementary material for "Characterization of sinking and suspended microeukaryotic communities in spring Oyashio waters": Fig. S1.pdf

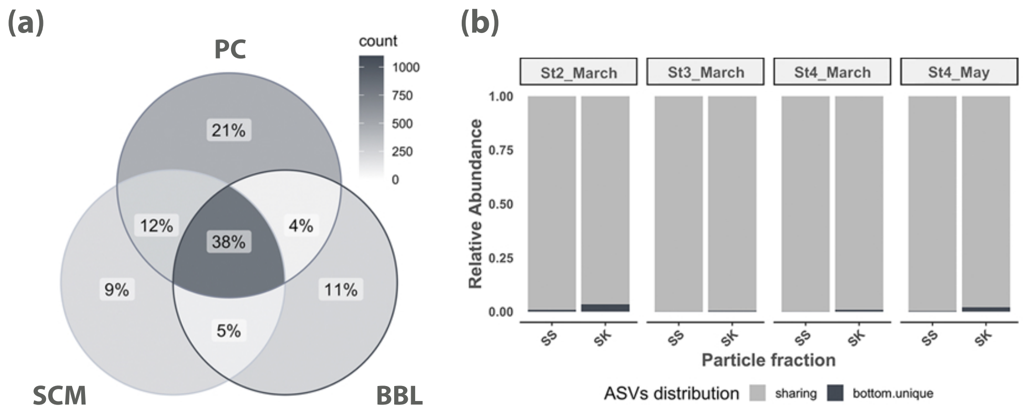


Fig. S1 Structural differences among the protist communities at the ASV level. (a) A Venn diagram representing the distribution of ASV richness across three depths; (b) The proportion of bottom unique ASVs in each bottom samples; X-axis indicate particle fractions: SS=Suspended particle, SK=Sinking Particle. Stations without bottom layer (St1 and St2 in May) were removed from the analysis.
