## Supplementary material for "Characterization of sinking and suspended microeukaryotic communities in spring Oyashio waters": Fig. S2.pdf

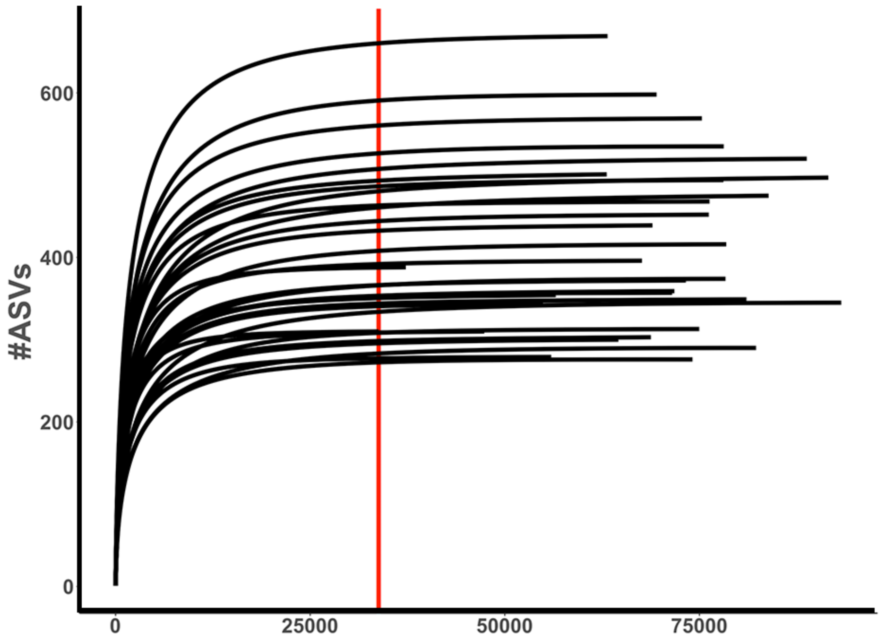


Fig. S2 Rarefaction curves of the ASV numbers for each sample. The vertical red line represents the smallest number of sequences per sample (33,794 reads).
