## Supplementary material for "Characterization of sinking and suspended microeukaryotic communities in spring Oyashio waters": Fig. S3.pdf

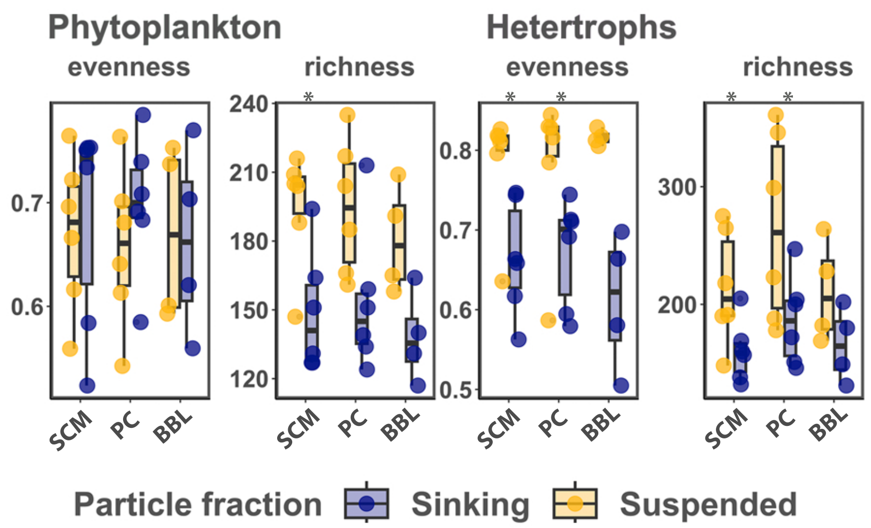


Fig. S3 Evenness and richness indices of different functional groups pooled by particle fractions and depths (SCM, PC, BBL). Stars indicate significant differences in indices between these two particle fractions, with * $p < 0.05$, ** $p < 0.01$, *** $p < 0.001$ (Wilcoxon test)
