## Supplementary material for "Characterization of sinking and suspended microeukaryotic communities in spring Oyashio waters": Fig. S4.pdf

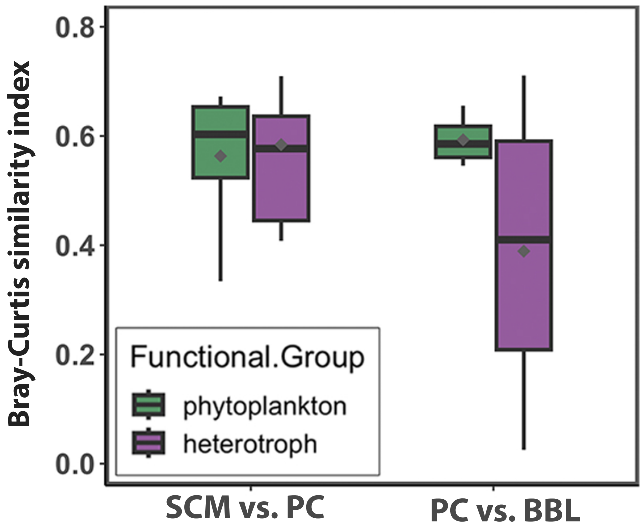


Fig. S4 Boxplot showing the Bray-Curtis similarity of suspended phytoplankton and heterotrophs between different depths. Outliers were removed from the boxplot. X-axis indicates depth pairs: SCM.vs.PC=suspended particles at SCM vs. suspended particles at PC, PC.vs.BBL=suspended particles at PC vs. suspended particles at BBL; Y-axis represents community similarity between depths for suspended particles; grey point represent the mean value of similarity.
