## Supplementary material for "Characterization of sinking and suspended microeukaryotic communities in spring Oyashio waters": Fig. S5.pdf

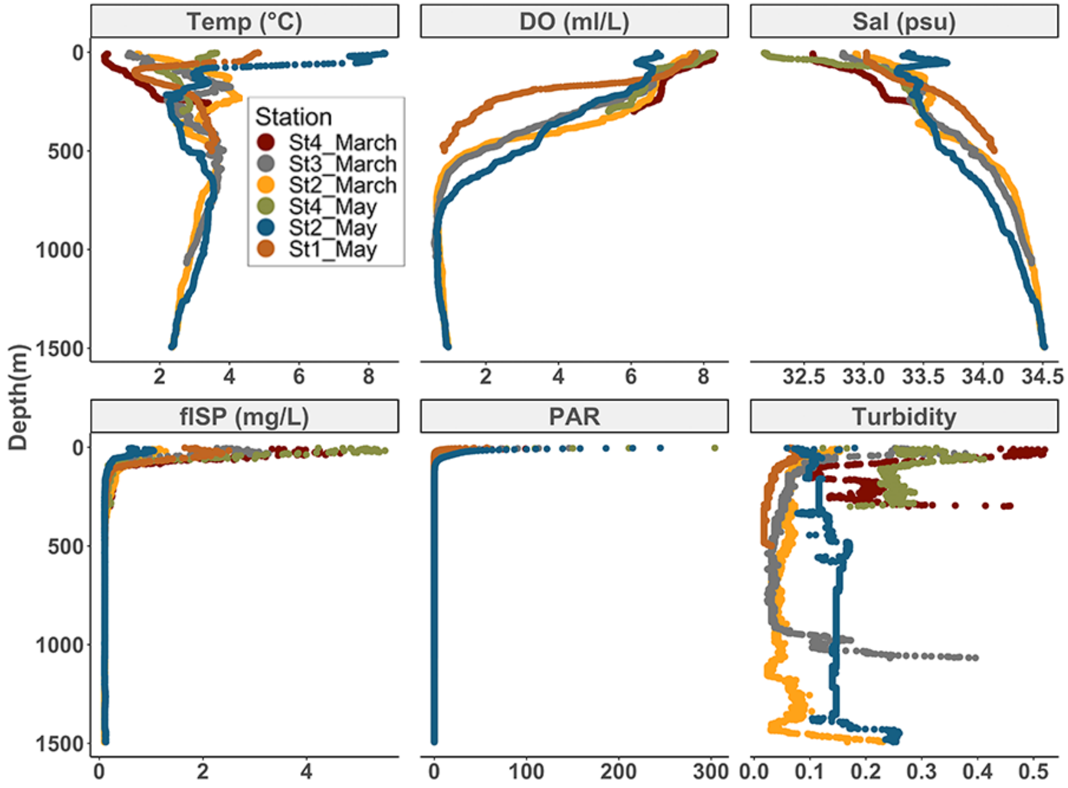


Fig. S5 Profiles of water column properties as measured using a CTD profiler. Sensors measured Temp, DO, Sal, chlorophyll fluorescence (flSP, mg/L), photosynthetically active radiation (PAR), and turbidity.
