## Supplementary material for "Characterization of sinking and suspended microeukaryotic communities in spring Oyashio waters": Table S1.docx

| **Sample ID** | **sites** | **Cruise** | **Layer^a^** | **Particle types^b^** | **# of raw pairs-end reads** | **# of raw ASVs** | **# of protists**  **ASV** | **# of phytoplankton ASVs** | **% of phytoplankton** | **# of heterotrophic**  **ASVs** | **% of heterotrophs** |
| --- | --- | --- | --- | --- | --- | --- | --- | --- | --- | --- | --- |
| A_M23_03_S | St2 | KS-21-4 | SCM | SS | 130,802 | 87567 | 524 | 208 | 87.41 | 316 | 12.59 |
| A_M22_03_F |  | (2021) |  | SK | 135,505 | 86438 | 432 | 195 | 78.07 | 237 | 21.93 |
| A_M24_03_S |  |  | PC | SS | 127,669 | 88087 | 558 | 225 | 86.88 | 333 | 13.12 |
| A_M19_03_F |  |  |  | SK | 119,037 | 76794 | 492 | 216 | 70.18 | 276 | 29.82 |
| A_M17_03_S |  |  | BBL | SS | 130,038 | 88101 | 466 | 216 | 83.55 | 250 | 16.45 |
| A_M18_03_F |  |  |  | SK | 136,236 | 85650 | 394 | 166 | 50.28 | 228 | 49.72 |
| A_M41_03_S | St3 |  | SCM | SS | 127,256 | 86752 | 456 | 226 | 80.99 | 230 | 19.01 |
| A_M39_03_F |  |  |  | SK | 150,900 | 100567 | 335 | 161 | 75.93 | 174 | 24.07 |
| A_M40_03_S |  |  | PC | SS | 121,070 | 83204 | 367 | 173 | 79.91 | 194 | 20.09 |
| A_M38_03_F |  |  |  | SK | 127,197 | 82518 | 309 | 154 | 71.27 | 155 | 28.73 |
| A_M36_03_S |  |  | BBL | SS | 127,303 | 85962 | 343 | 174 | 83.69 | 169 | 16.31 |
| A_M37_03_F |  |  |  | SK | 121,331 | 79296 | 309 | 132 | 51.06 | 177 | 48.94 |
| A_M15_03_S | St4 |  | SCM | SS | 103,290 | 71928 | 347 | 151 | 55.48 | 196 | 44.52 |
| A_M13_03_F |  |  |  | SK | 124,156 | 88588 | 284 | 143 | 49.41 | 141 | 50.59 |
| A_M14_03_S |  |  | PC | SS | 104,702 | 75012 | 351 | 173 | 57.70 | 178 | 42.30 |
| A_M12_03_F |  |  |  | SK | 104,583 | 78101 | 296 | 136 | 55.26 | 160 | 44.74 |
| A_M10_03_S |  |  | BBL | SS | 118,297 | 77025 | 354 | 168 | 67.50 | 186 | 32.50 |
| A_M11_03_F |  |  |  | SK | 105,405 | 77489 | 273 | 118 | 46.22 | 155 | 53.78 |
| A_M06_05_S | St1 | KS-21-7 | SCM | SS | 125,359 | 86177 | 410 | 193 | 57.35 | 217 | 42.65 |
| A_M06_05_F |  | (2021) |  | SK | 121,499 | 80630 | 344 | 165 | 49.00 | 179 | 51.00 |
| A_M05_05_S |  |  | PC | SS | 135,582 | 93213 | 437 | 208 | 56.09 | 229 | 43.91 |
| A_M05_05_F |  |  |  | SK | 105,360 | 68752 | 338 | 142 | 61.13 | 196 | 38.87 |
| A_M12_05_S | St2 |  | SCM | SS | 128,592 | 83554 | 438 | 211 | 57.16 | 227 | 42.84 |
| A_M12_05_F |  |  |  | SK | 122,149 | 77926 | 277 | 127 | 35.67 | 150 | 64.33 |
| A_M11_05_S |  |  | PC | SS | 126,013 | 85144 | 661 | 238 | 48.78 | 423 | 51.22 |
| A_M11_05_F |  |  |  | SK | 118,060 | 77286 | 388 | 159 | 40.66 | 229 | 59.94 |
| A_M21_05_S | St4 |  | SCM | SS | 140,181 | 93007 | 500 | 207 | 62.95 | 293 | 37.05 |
| A_M21_05_F |  |  |  | SK | 115,273 | 57547 | 310 | 132 | 56.34 | 178 | 43.66 |
| A_M20_05_S |  |  | PC | SS | 109,736 | 71533 | 592 | 197 | 53.91 | 395 | 46.09 |
| A_M20_05_F |  |  |  | SK | 118,150 | 85612 | 369 | 124 | 32.63 | 245 | 67.37 |
| A_M19_05_S |  |  | BBL | SS | 124,422 | 83227 | 488 | 200 | 61.94 | 288 | 38.06 |
| A_M19_05_F |  |  |  | SK | 108,510 | 74420 | 352 | 141 | 47.12 | 211 | 52.88 |

Table S1. Overview of the 18S metabarcoding data used in this study

^a^ SCM: Subsurface Chlorophyll Maximum; PC: Pycnocline; BBL: Bottom Boundary Layer

^b^ SS: Suspended; SK: Sinking
