## Supplementary material for "Characterization of sinking and suspended microeukaryotic communities in spring Oyashio waters": Table S2.docx

Table S2. Number of unique ASVs within taxonomic groups that are either photosynthetic or have the potential for photosynthesis as defined by the PR2 database (except dinoflagellates)

| **Phytoplankton Taxa** | |  |  |  |
| --- | --- | --- | --- | --- |
| **Kingdom (PR2 Level1)** | **Supergroup (PR2 level2)** | **Division/Phylumn (PR2 level3)** | **Class (PR2 level4)** | **# of ASVs** |
| Eukaryota | Archaeplastida | Chlorophyta | Chlorophyceae | 2 |
| Eukaryota | Archaeplastida | Chlorophyta | Pyramimonadophyceae | 22 |
| Eukaryota | Archaeplastida | Chlorophyta | Trebouxiophyceae | 2 |
| Eukaryota | Archaeplastida | Chlorophyta | Mamiellophyceae | 14 |
| Eukaryota | Hacrobia | Cryptophyta | Cryptophyceae | 27 |
| Eukaryota | Hacrobia | Haptophyta | Haptophyta_Clade_HAP3 | 3 |
| Eukaryota | Hacrobia | Haptophyta | Haptophyta_Clade_HAP4 | 3 |
| Eukaryota | Hacrobia | Haptophyta | Rappephyceae | 4 |
| Eukaryota | Hacrobia | Haptophyta | Prymnesiophyceae | 87 |
| Eukaryota | Rhizaria | Cercozoa | Chlorarachniophyceae | 10 |
| Eukaryota | Stramenopiles | Ochrophyta | Bacillariophyta | 144 |
| Eukaryota | Stramenopiles | Ochrophyta | Bolidophyceae | 7 |
| Eukaryota | Stramenopiles | Ochrophyta | Chrysophyceae | 29 |
| Eukaryota | Stramenopiles | Ochrophyta | Dictyochophyceae | 1 |
| Eukaryota | Stramenopiles | Ochrophyta | MOCH-2 | 3 |
| Eukaryote | Stramenopiles | Ochrophyta | MOCH-3 | 1 |
| Eukaryota | Stramenopiles | Ochrophyta | Pelagophyceae | 7 |
| Eukaryota | Stramenopiles | Ochrophyta | Raphidophyceae | 2 |
| Total |  |  |  | 368 |
