## Supplementary material for "Characterization of sinking and suspended microeukaryotic communities in spring Oyashio waters": Table S3.docx

Table S3. Dinoflagellate ASVs that have the potential for autotrophy or heterotrophy, based on current studies.

| **Class (PR2 level4)** | **Order (PR2 level5)** | **Family (PR2 level6)** | **Genus (PR2 level7)** | **Species (PR2 level8)** | **Functional Group^a^** | **Reference Source** |
| --- | --- | --- | --- | --- | --- | --- |
| Dinophyceae | Dinophyceae_X | Dinophyceae_XX | Abedinium | Abedinium_dasypus | Heterotroph | (Cooney, 2022) |
| Dinophyceae | Dinophyceae_X | Dinophyceae_XX | Dinophyceae_XXX | Dinophyceae_XXX_sp. | Phytoplankton | –– |
| Dinophyceae | Dinophysiales | Dinophysiaceae | Dinophysis | unassigned | Phytoplankton | (Reguera et al., 2012) |
| Dinophyceae | Dinophysiales | Dinophysiaceae | Ornithocercus | unassigned | Heterotroph | (Kim et al., 2021) |
| Dinophyceae | Dinophysiales | Dinophysiaceae | unassigned | unassigned | Heterotroph | (Kim et al., 2021) |
| Dinophyceae | Dinophysiales | Oxyphysiaceae | Phalachroma | Phalacroma_rotundatum | Phytoplankton | (Caroppo et al., 1999) |
| Dinophyceae | Dinophysiales | Oxyphysiaceae | Phalachroma | unassigned | Phytoplankton | (Caroppo et al., 1999) |
| Dinophyceae | Gonyaulacales | Ceratiaceae | Tripos | Tripos_fusus | Phytoplankton | (Hallegraeff et al., 2020) |
| Dinophyceae | Gonyaulacales | Ceratiaceae | Tripos | unassigned | Phytoplankton | (Hallegraeff et al., 2020) |
| Dinophyceae | Gonyaulacales | Goniodomataceae | Alexandrium | Alexandrium_fundyense | Phytoplankton | (Anderson et al., 2005) |
| Dinophyceae | Gonyaulacales | Gonyaulacaceae | Amylax | Amylax_buxus | Phytoplankton | (Koike and Takishita, 2008) |
| Dinophyceae | Gonyaulacales | Gonyaulacaceae | Gonyaulax | Gonyaulax_spinifera | Phytoplankton | (Rhodes et al., 2006) |
| Dinophyceae | Gonyaulacales | Protoceratiaceae | Ceratocorys | Ceratocorys_horrida | Phytoplankton | (Zirbel et al., 2000) |
| Dinophyceae | Gonyaulacales | Pyrophacaceae | Fragilidium | –– | Phytoplankton | (Skovgaard et al., 2000) |
| Dinophyceae | Gymnodiniales | Chytriodiniaceae | Chytriodinium | Chytriodinium_roseum | Heterotroph | (GÓmez et al., 2009) |
| Dinophyceae | Gymnodiniales | Gymnodiniaceae | Balechina | Balechina_pachydermata | Heterotroph | (Gómez et al., 2015) |
| Dinophyceae | Gymnodiniales | Gymnodiniaceae | Gymnodinium | Gymnodinium_dorsalisulcum | Heterotroph | (Murray et al., 2007) |
| Dinophyceae | Gymnodiniales | Gymnodiniaceae | Gyrodinium | Gyrodinium_dominans | Heterotroph | (Jae and Hae, 2004) |
| Dinophyceae | Gymnodiniales | Gymnodiniaceae | Gyrodinium | Gyrodinium_helveticum | Heterotroph | (Takano and Horiguchi, 2004) |
| Dinophyceae | Gymnodiniales | Gymnodiniaceae | Gyrodinium | Gyrodinium_heterogrammum | Heterotroph | (Larsen, 1994) |
| Dinophyceae | Gymnodiniales | Gymnodiniaceae | Gyrodinium | Gyrodinium_rubrum | Heterotroph | (Takano and Horiguchi, 2004) |
| Dinophyceae | Gymnodiniales | Gymnodiniaceae | Gymnodinium | –– | Phytoplankton | (Seki et al., 1995; Carreto et al., 2006) |
| Dinophyceae | Gymnodiniales | Kareniaceae | –– | –– | Phytoplankton | (Bergholtz et al., 2006) |
| Dinophyceae | Gymnodiniales | Warnowiaceae | Warnowia | Warnowia_sp. | Heterotroph | (GÓmez et al., 2009) |
| Dinophyceae | Gymnodiniales | unassigned | unassigned | unassigned | Phytoplankton | (Seki et al., 1995; Carreto et al., 2006) |
| Dinophyceae | Peridiniales | Amphidiniopsidaceae | –– | –– | Heterotroph | (Head et al., 2001; Gomez, 2012) |
| Dinophyceae | Peridiniales | Amphidomataceae | Azadinium | unassigned | Phytoplankton | (Jauffrais et al., 2012) |
| Dinophyceae | Peridiniales | Blastodiniaceae | Blastodinium | Blastodinium_spinulosum | Heterotroph | (Skovgaard and Salomonsen, 2009) |
| Dinophyceae | Peridiniales | Blastodiniaceae | Blastodinium | unassigned | Heterotroph | (Coats et al., 2008) |
| Dinophyceae | Peridiniales | Diplopsalidaceae | Diplopsalis | unassigned | Phytoplankton | (Reguera et al., 2012) |
| Dinophyceae | Peridiniales | Diplopsalidaceae | Niea | Niea_acanthocysta | Heterotroph | (Matsuoka et al., 2018) |
| Dinophyceae | Peridiniales | Heterocapsaceae | Heterocapsa | –– | Phytoplankton | (Iwataki, 2008) |
| Dinophyceae | Peridiniales | Protoperidiniaceae | Protoperidinium | –– | Heterotroph | (Olenina et al.,2006; Jeong et al., 2010) |
| Dinophyceae | Peridiniales | Thoracosphaeraceae | –– | –– | Phytoplankton | (Gottschling et al., 2020) |
| Dinophyceae | Prorocentrales | Prorocentraceae | Prorocentrum | –– | Phytoplankton | (Hernández-Becerril et al., 2000) |
| Dinophyceae | Suessiales | Suessiaceae | Biecheleria | unassigned | Phytoplankton | (Jang et al., 2015) |
| Dinophyceae | Suessiales | Suessiales_X | Dactylodinium | Dactylodinium_pterobelotum | Phytoplankton | (Takahashi et al., 2017) |
| Dinophyceae | Torodiniales | Torodiniaceae | Torodinium | unassigned | Phytoplankton | (Larsen, 1994) |
| Ellobiophyceae | Thalassomycetales | Thalassomycetaceae | Ellobiopsis | Ellobiopsis_chattonii | Heterotroph | (Horiguchi, 2015) |
| Syndiniales | –– | –– | –– | –– | Heterotroph | –– |

^a^ Mixtrophs were simply grouped into phytoplankton

Olenina, I., Hajdu, S., Edler, L., Andersson, A., Wasmund, N., Busch, S., Göbel, J., Gromisz, S., Huseby, S., Huttunen, M.Jaanus, A., Kokkonen, P., Ledaine, I. and Niemkiewicz, E. Biovolumes and Size-Classes of Phytoplankton in the Baltic Sea Helsinki Commission Baltic Marine Environment Protection Commission.

Reguera, B., Velo-Suárez, L., Raine, R., and Park, M.G. (2012) Harmful Dinophysis species: A review. *Harmful Algae* **14**: 87–106.

Rhodes, L., McNabb, P., De Salas, M., Briggs, L., Beuzenberg, V., and Gladstone, M. (2006) Yessotoxin production by Gonyaulax spinifera. *Harmful Algae* **5**: 148–155.

Seki, T., Satake, M., Mackenzie, L., Kaspar, H.F., and Yasumoto, T. (1995) Gymnodimine, a new marine toxin of unprecedented structure isolated from New Zealand oysters and the dinoflagellate, Gymnodinium sp. *Tetrahedron Lett* **36**: 7093–7096.

Skovgaard, A., Hansen, P.J., and Stoecker, D.K. (2000) Physiology of the mixotrophic dinoflagellate Fragilidium subglobosum. I. Effects of phagotrophy and irradiance on photosynthesis and carbon content. *Mar Ecol Prog Ser* **201**: 129–136.

Skovgaard, A. and Salomonsen, X.M. (2009) Blastodinium galatheanum sp. nov. (Dinophyceae) a parasite of the planktonic copepod Acartia negligens (Crustacea, Calanoida) in the central Atlantic Ocean. *Eur J Phycol* **44**: 425–438.

Takahashi, K., Moestrup, Ø., Wada, M., Ishimatsu, A., Nguyen, V.N., Fukuyo, Y., and Iwataki, M. (2017) Dactylodinium pterobelotum gen. et sp. nov., a new marine woloszynskioid dinoflagellate positioned between the two families Borghiellaceae and Suessiaceae. *J Phycol* **53**: 1223–1240.

Takano, Y. and Horiguchi, T. (2004) Surface ultrastructure and molecular phylogenetics of four unarmored heterotrophic dinoflagellates, including the type species of the genus Gyrodinium (Dinophyceae). *Phycological Res* **52**: 107–116.

Zirbel, M.J., Veron, F., and Latz, M.I. (2000) THE REVERSIBLE EFFECT OF FLOW ON THE MORPHOLOGY OF CERATOCORYS HORRIDA Most cells experience an active and variable fluid environment , in which hydrodynamic forces can af- fect aspects of cell physiology including gene regula- tion , growth , nutrient up. **58**: 46–58.
