## Supplementary material for "Characterization of sinking and suspended microeukaryotic communities in spring Oyashio waters": Table S4.docx

Table S4. Environmental variables of all samples in the Oyashio waters

| Sites | Cruise | Date | Long. | Lat. | Seafloor  depth (m) | Chl a (µg/L) | MSC Type^a^ | Layer | Depth (m) | Temp (°C) | Sal (‰) | DO  (l/L) | NO_3_^b^ | NO_2_^b^ | NH_4_^b^ | PO_4_^b^ | SiOH_4_^b^ | POC flux^c^ | PON flux^c^ |
| --- | --- | --- | --- | --- | --- | --- | --- | --- | --- | --- | --- | --- | --- | --- | --- | --- | --- | --- | --- |
| St2 | KS-21-4 | 15^th^ Mar | 142.9 | 41.4 | 1508 | 1.17 | small | SCM | 17 | 1.37 | 32.93 | 7.62 | 17.94 | 0.24 | < 0.1^d^ | 1.50 | 33.39 | 89.69 | 18.36 |
|  | (2021) |  |  |  |  |  | small | PC | 150 | 3.93 | 33.51 | 6.68 | 19.94 | 0.18 | < 0.1 | 1.56 | 37.31 | 0 | 0 |
|  |  |  |  |  |  |  | giant | BBL | 1478 | 2.40 | 34.50 | 0.95 | 43.87 | 0.06 | < 0.1 | 3.13 | 163.23 | 30.94 | 4.49 |
| St3 |  | 18^th^ Mar | 142.7 | 41.8 | 1078 | 5.4 | small | SCM | 30 | 1.63 | 32.89 | 7.93 | 14.63 | 0.32 | < 0.1 | 1.34 | 29.34 | 153.40 | 30.84 |
|  |  |  |  |  |  |  | small | PC | 250 | 2.22 | 33.41 | 5.44 | 28.05 | 0.08 | < 0.1 | 2.17 | 58.71 | 0 | 0 |
|  |  |  |  |  |  |  | giant | BBL | 1050 | 2.80 | 34.41 | 0.70 | 43.85 | 0.09 | < 0.1 | 3.19 | 155.52 | 29.77 | 3.74 |
| St4 |  | 14^th^ Mar | 142.7 | 42.0 | 310 | 11.93 | small | SCM | 11 | 0.47 | 32.57 | 8.28 | 9.60 | 0.18 | < 0.1 | 1.08 | 23.39 | 228.71 | 49.71 |
|  |  |  |  |  |  |  | small | PC | 71 | 0.63 | 32.79 | 7.60 | 15.49 | 0.18 | < 0.1 | 1.40 | 29.99 | 717.60 | 57.76 |
|  |  |  |  |  |  |  | giant | BBL | 289 | 1.76 | 33.47 | 6.12 | 19.32 | 0.23 | < 0.1 | 1.62 | 37.47 | 86.96 | 12.74 |
| St1 | KS-21-7 | 8^th^ May | 144.0 | 41.0 | 4357 | 1.51 | small | SCM | 30 | 2.19 | 33.02 | 7.80 | 9.30 | 0.16 | 1.05 | 1.03 | 2.95 | 385.34 | 64.79 |
|  | (2021) |  |  |  |  |  | small | PC | 65 | 4.54 | 33.09 | 7.20 | 23.73 | 0.40 | 1.47 | 2.03 | 41.12 | 274.79 | 53.54 |
| St2 |  | 9^th^ May | 142.9 | 41.4 | 1518 | 1.36 | small | SCM | 14 | 8.11 | 33.40 | 6.80 | 2.60 | 0.09 | < 0.1 | 0.33 | 4.55 | 152.74 | 32.01 |
|  |  |  |  |  |  |  | small | PC | 65 | 5.52 | 33.49 | 6.31 | 12.03 | 0.19 | 0.56 | 1.04 | 17.95 | 103.29 | 22.10 |
| St4 |  | 6^th^ May | 142.7 | 42.0 | 318 | 7.94 | small | SCM | 25 | 3.30 | 32.25 | 8.01 | 3.34 | 0.20 | 0.62 | 0.62 | 2.15 | 755.68 | 129.66 |
|  |  |  |  |  |  |  | small | PC | 90 | 1.93 | 33.18 | 6.69 | 20.48 | 0.27 | 0.63 | 1.77 | 36.57 | 208.48 | 39.90 |
|  |  |  |  |  |  |  | giant | BBL | 305 | 2.83 | 33.18 | 5.37 | 27.38 | 0.22 | < 0.1 | 2.11 | 54.42 | 352.65 | 63.65 |

^a^ Small MSC: upper 100L, base 20L ; Giant MSC: upper 300L, base 70 L

^b^ Macronutrient unit: µmol/L

^c^ PON and POC flux: mg/m2/d

^d^ lower than detection limit (0.1 µmol/L)
