## Supplementary material for "Characterization of sinking and suspended microeukaryotic communities in spring Oyashio waters": Table S5.docx

Table S5. PERMANOVA results

| Source | Df^a^ | SumOfSqs^b^ | R2^c^ | F^d^ | Pr(>F)^e^ |
| --- | --- | --- | --- | --- | --- |
| As.factor(particle types:suspended & sinking) | 1 | 0.5265432 | 0.09603109 | 3.186982 | 0.003 |
| Residual | 30 | 4.9565055 | 0.90396891 | NA | NA |
| Total | 31 | 5.4830487 | 1.00000000 | NA | NA |
| As.factor(condition:diatom-dominant & dinoflagellate-abundant) | 1 | 1.372062 | 0.2502371 | 10.01265 | 1e-04 |
| Residual | 30 | 4.110986 | 0.7497629 | NA | NA |
| Total | 31 | 5.483049 | 1.0000000 | NA | NA |

^a^ Df: stands for degrees of freedom

^b^ SumOfSqs: stands for sum of squares

^c^ R2: coefficient of determination

^d^ F: F-statistic

^e^ Pr(>F): *p*-value associated with the *F*-statistic
